# The molecular basis of sugar detection by an insect taste receptor

**DOI:** 10.1101/2023.12.18.572227

**Authors:** João Victor T. Gomes, Shivinder Singh-Bhagania, Matthew Cenci, Carlos Chacon Cordon, Manjodh Singh, Joel A. Butterwick

## Abstract

Animals crave sugars because of their energy potential and the pleasurable sensation of tasting sweetness. Yet all sugars are not metabolically equivalent, requiring mechanisms to detect and differentiate between chemically similar sweet substances. Insects use a family of ionotropic gustatory receptors to discriminate sugars, each of which is selectively activated by specific sweet molecules. To gain insight into the molecular basis of sugar selectivity, we determined structures of Gr9, a gustatory receptor from the silkworm Bombyx mori (BmGr9), in the absence and presence of its sole activating ligand, D-fructose. These structures, along with structure-guided mutagenesis and functional assays, illustrate how specificity for D-fructose is seemingly achieved by a ligand-binding pocket that precisely matches the overall shape and pattern of chemical groups in D-fructose. However, our computational docking and experimental binding assays revealed that other sugars also bind BmGr9, yet they are unable to activate the receptor. We identified the conformational change required to open the channel gate that provides an additional layer of receptor tuning in BmGr9; only D-fructose can both fit into the pocket and simultaneously engage a bridge of two conserved aromatic residues that connects the pocket to the ion conducting pore. Thus, chemical specificity does not depend solely on the selectivity of the ligand-binding pocket, but it is an emergent property arising from a combination of receptor-ligand interactions and allosteric coupling. Our results support a model whereby coarse receptor tuning is derived from the size and chemical characteristics of the pocket, whereas fine-tuning of receptor activation is achieved through the selective engagement of an allosteric pathway that regulates ion conduction.

---

Sugars are a primary source of energy and influence nutrient sensing, metabolic responses, reward mechanisms, and taste perception^1,2^. Most animals can taste sugars via sweet receptors expressed in dedicated taste organs^3^. Mammals taste all sweet compounds using a single heterodimeric taste receptor, composed of two G-protein-coupled receptors (T1R2 and T1R3)^4^. Insects, instead, rely on a distinct family of sweet gustatory receptors (GRs)^5^, which are tetrameric ligand-gated cation channels^6^. A distinctive feature of these GRs is that each receptor specializes in detecting only a subset of sugar molecules^6– 10^, despite the high degree of chemical similarity among saccharides. An extreme example of sugar specialization occurs in a conserved subfamily of GRs founded by *Drosophila melanogaster* Gr43a (DmGr43a), whose members are selectively activated only by the monosaccharide D-fructose^6,7^, providing a unique opportunity to investigate how specificity for a single sugar is achieved.

Here, we investigate the structural basis of sugar discrimination by Gr9 (BmGr9), the silk moth *Bombyx mori* ortholog of DmGr43a. We determined structures of BmGr9 in multiple gating states: closed, in the absence of sugar, and opened, in the presence of D-fructose. BmGr9 harbours a sugar-binding pocket in the transmembrane region of each subunit that tightly envelopes D-fructose and precisely coordinates its hydroxyl groups via a set of conserved polar amino acids. Despite these seemingly specific interactions, computational docking and experimental binding studies suggest that other sugar molecules also fit into the BmGr9 pocket, but they are unable to activate the receptor. Thus, the geometric arrangement of chemical groups within the ligand-binding pocket does not appear to be sufficient to explain the selective activation of this receptor by only D-fructose. We find that D-fructose is unique in its ability to induce a conformational change that is required to open the channel gate. Activation efficacy, therefore, depends on residues that extend beyond the receptor-ligand interactions that occur in the binding pocket. Our findings show how narrow chemical tuning can be achieved in taste receptors and suggests how the tuning of chemoreceptors could be adjusted to recognize different regions of chemical space.

## Structure of BmGr9 bound to D-fructose

We confirmed the narrow chemical tuning of BmGr9 by transiently co-expressing the receptor with a fluorescent calcium reporter, GCaMP6s^11^, in HEK293 cells and recorded fluorescence changes upon addition of sweet compounds. Of a panel of 27 naturally occurring sugars and artificial sweeteners, we found that only D-fructose elicited a strong response (**Fig. 1A; Sup. Fig. 1**), which is consistent with previous results^6^. Fitting the Hill equation to dose-response data yielded a half-maximal activation concentration (*EC*_50_) of 8 mM for D-fructose (**Fig. 1B**). Such low affinities are common for sugar-sensing GRs^6,9^ and likely reflect the high concentrations of sugars typically found in floral nectars^12^ and insect hemolymph^7^. Notably, even sugars that are highly structurally similar to D-fructose did not activate BmGr9 significantly, such as L-sorbose, an epimer of fructose differing only by the relative orientation of a single hydroxyl group (**Fig. 1A,B**).

**Figure 1:**
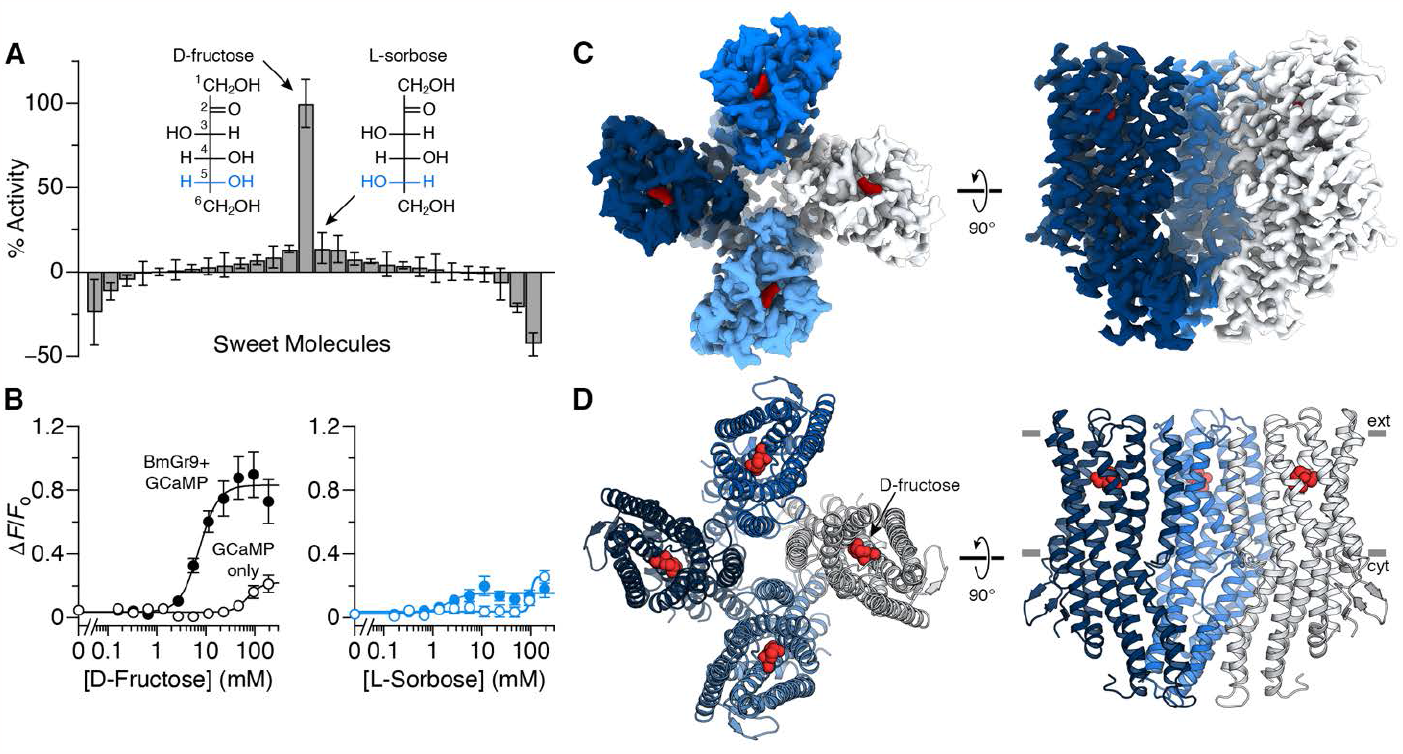
Structure of an insect taste receptor. (**A**) Activation of BmGr9 by a panel of sweet compounds, measured as the change in fluorescence relative to D-fructose (*n* = 4, bars are mean ± s.e.m.). Insets show Fischer projections of epimers D-fructose and L-sorbose, with carbons numbered and difference highlighted in blue. (**B**) Dose-response of raw fluorescence changes of HEK293 cells transfected with BmGr9 and GCaMP (closed circles) or GCaMP only (open circles) when titrated with D-fructose (left) or L-sorbose (right). D-Fructose data are best fit by an *EC*_50_ of 8.2 (5.9–11.5) mM with a Hill coefficient of 2.3 (1.3–4.6) (*n* = 8; points are mean ± s.e.m.; fitted 95% confidence intervals are given in parentheses). For L-sorbose (*n* = 5), the maximum activity is less than control wells with GCaMP only. (**C,D**) Cryo-EM density map (**C**) and ribbon model (**D**) of BmGr9 bound to D-fructose (red) shown from the top (left) and side (right). Approximate boundaries for the extracellular (ext) and cytoplasmic (cyt) sides are indicated. In (**C**) and (**D**), the front subunit has been removed from the side views to expose the pore.

To investigate the structural basis for the remarkable sugar specificity of BmGr9, we purified the receptor in the presence of a saturating amount of D-fructose (**Sup. Fig. 2**) and determined the structure of the complex using single-particle cryogenic electron microscopy (cryo-EM). Three-dimensional reconstruction with four-fold averaging yielded a density map with 3.0 Å overall resolution (**Fig. 1C; Sup. Fig. 3A-D**), which allowed us to build a model for the majority of the protein (**Fig. 1D**). The structure of BmGr9 closely resembles insect olfactory receptors (ORs)^13,14^, demonstrating how the same overall architecture underlies detection of tastants and odorants in insects, unlike mammalian receptors that recognize these compounds through distinct families using different binding domains^15^.

### Sugar-binding pocket

D-Fructose is predicted to bind within an extracellular-facing pocket formed by the S1-S6 transmembrane helices of each BmGr9 subunit^16^. We observed additional density in the putative sugar-binding pocket, not attributable to protein, that is the approximate size and shape of a monosaccharide. In aqueous solution, D-fructose rapidly interconverts between five-membered (α- and β-furanose) and six-membered (β-pyranose) ring configurations^17,18^. To help determine which form of D-fructose is bound in our structure, we computationally docked fructose conformers into the pocket using AutoDock Vina^19,20^. We found β-furanose and β-pyranose forms docked with the lowest energy scores (−5.7 kcal/mol), while the calculated energetics of α-furanose was less favourable (−5.0 kcal/mol; **Sup. Fig. 4A-C)**. We therefore refined two models of BmGr9, bound to β-D-fructopyranose or β-D-fructofuranose, which yielded nearly identical fructose conformations (**Sup. Fig. 4D-F**). In what follows, we focus on β-D-fructopyranose, since it is the major conformer (∼75%)^17,18^ and thus most likely to be bound by BmGr9, but both forms appear capable of making similar interactions with BmGr9.

The sugar-binding pocket in BmGr9 extends from the extracellular surface to almost halfway through the membrane. D-Fructose sits at the base of this pocket, approximately 15 Å from the extracellular surface, making direct contact with residues in helices S2-S6 (**Fig. 2A,B**). D-Fructose is oriented such that its hydrophobic and hydrophilic surfaces make very different interactions with residues within the binding pocket. Notably, one of the hydrophobic faces of β-D-fructopyranose, formed by hydrogens on C1 and C3, borders Trp354 in S6 at the side of the pocket (**Fig. 2C**). In protein regions that interact with sugar molecules, tryptophan and other aromatic residues are common due to the favourable formation of CH–π interactions^21^. The distinct hydrophobic regions of each sugar dictate the preferred orientation of the molecule when bound to its cognate receptor^21^. Thus, the Trp-fructose interaction likely positions the sugar within the binding pocket.

**Figure 2:**
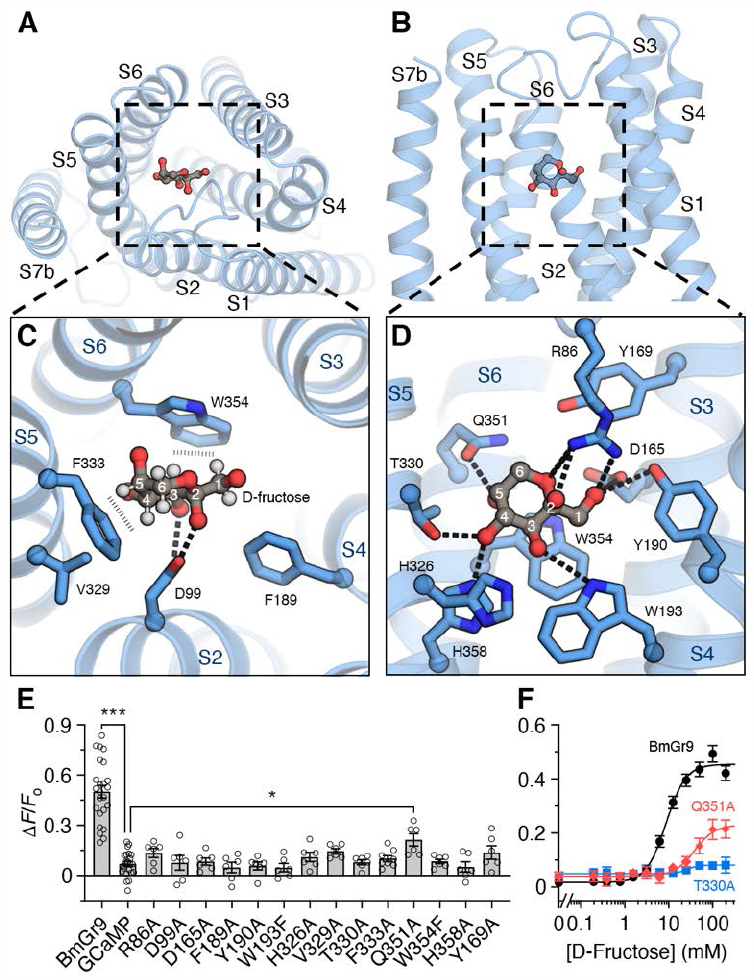
Sugar-binding pocket in BmGr9. (**A,B**) Position of β-D-fructopyranose (grey carbons and red oxygens) within a subunit of BmGr9 (blue) viewed from the top (**A**) and side (**B**). (**C,D**) Close-up views showing interactions between BmGr9 and D-fructose (with carbons numbered). Hydrogen-bonding interactions are drawn as dashed lines. In (**C**), hydrogens on D-fructose (white) are shown to highlight hydrophobic interactions with Trp354 and Phe333 (indicated by vertical dashes). (**E**) Effect of mutations within the sugar-binding pocket on the activity of BmGr9. Bars are mean ± s.e.m with individual replicates shown as open circles. Only wild-type BmGr9 (*P* < 0.0001) and Q351A (*P* < 0.05) have significantly different activity than GCaMP alone. (**F**) Dose response of select mutants from (**E**). Data points are mean ± s.e.m from *n* = 6–12 experiments.

The hydroxyl groups in D-fructose are all coordinated by polar groups (**Fig. 2C,D**), from acidic (Asp99, Asp165), basic (Arg86, His358), and uncharged residues (Tyr190, Trp193, Thr330, and Gln351). Individual hydroxyls often interact with multiple amino acids, creating a network of bridges between transmembrane helices. For example, hydroxyl groups on C1 (bridging S3 and S4), C3 (S2 and S4), and C4 (S5 and S6) all link neighbouring helices, likely imparting considerable stability to the pocket when sugar is present.

Amino acids lining the pocket are highly conserved among fructose-selective receptors, but not in other sweet-sensing GRs (**Sup. Fig. 5**), providing a rationale for the differing sugar sensitivities among GRs. Mutation of the aromatic and polar residues within the pocket to Ala (or Trp to Phe) eliminated activation by D-fructose, with the exception of Gln351 to Ala, which retained only marginal activity (**Fig. 2E,F**). More conservative mutations were also generally not tolerated. For instance, mutating either Asp99 or Asp165 to Asn, Gln, or Glu resulted in non-functional channels, but Arg86 to Lys retained activity (**Sup. Fig. 6**). The strict requirement for specific amino acids suggests that their distribution within the binding pocket of BmGr9 forms a precise geometric arrangement to coordinate D-fructose.

### Conserved chemoreceptor ligand-binding locus

Comparing the structure of fructose-bound BmGr9 to an eugenol-bound OR from the jumping bristletail *Machilis hrabei* (MhOr5)^14^ shows that sugar and odorant occupy similar locations (**Fig. 3A,B**), suggesting that the ligand-binding locus is conserved across the insect chemoreceptor superfamily. However, BmGr9 and MhOr5 contrast in several critical aspects. Notably, sugar and odorants occupy different regions of the pocket. D-Fructose sits close to the pore at the inner edge of the pocket, partially exposed to the extracellular solution and interacting closely with many residues along S5 and S6 (**Fig. 3C,D**). Eugenol, instead, sits in an occluded cavity adjacent to S3 at the outer edge, approximately 6 Å distal and 6 Å deeper than fructose, too far to directly interact with S5. In the presence or absence of an odorant, the ligand-binding pocket of MhOr5 is enclosed by protein (**Fig. 3E**)^14^. Hydrophobic odorants have been suggested to enter the binding pocket from the membrane through a tunnel formed transiently between S3 and S6, which would allow odorants to access the pocket near the outer edge^22^. Since water-soluble sugars appear to enter the pocket directly from the extracellular space, the positional differences of the ligands observed here may represent a distinctive feature of GRs versus ORs.

**Figure 3:**
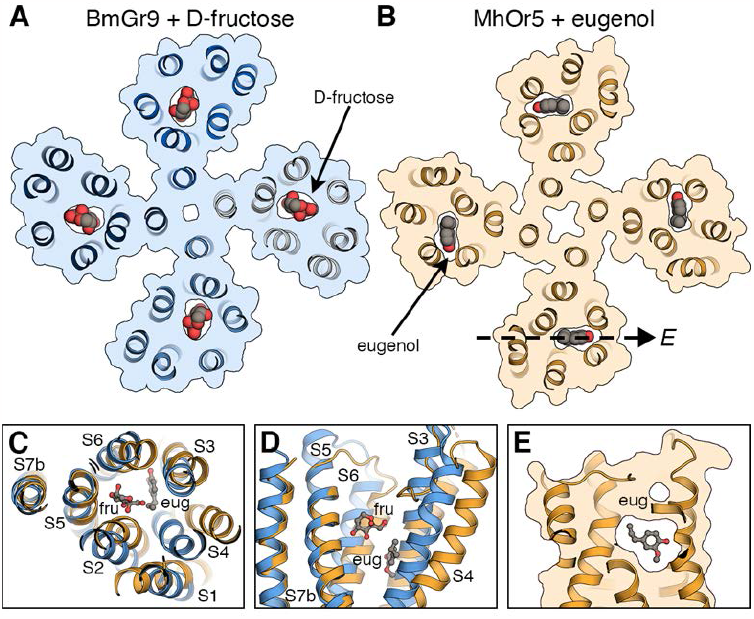
Ligand-binding locus is conserved among insect chemoreceptors. (**A,B**) Slice through the pockets of BmGr9 bound to D-fructose (**A**) and MhOr5 bound to eugenol (**B**; PDB 7LID) highlighting the positions of their respective ligands. (**C,D**) Superposition of a single subunit of BmGr9 (blue) and MhOr5 (orange) showing the relative positions of fructose (fru) and eugenol (eug), viewed from the top (**C**) and side (**D**). (**E**) Vertical slice through the pocket of MhOr5, in a similar orientation as Fig. 5B,C. Eugenol is encapsulated within MhOr5 with no direct means of egress.

The structural and chemical characteristics of the ligand-binding pockets are also different between BmGr9 and MhOr5. BmGr9 binds its single ligand with a hydrophilic pocket built to recognize the specific molecular features of D-fructose, whereas MhOr5 engages diverse odorants with a promiscuous hydrophobic pocket that accommodates differently shaped molecules^14^, consistent with the vastly different tuning profiles of these two chemoreceptors.

### Gating and cooperativity among subunits

To gain insight into the mechanism of receptor activation, we also determined the structure of BmGr9 in the absence of ligand. We resolved a similar quality map as the fructose-bound BmGr9 (**Sup. Fig. 3E-H**) but with the extracellular hydrophobic gate closed, consistent with the absence of any density in the ligand-binding pocket and representing a closed state (**Fig. 4A,B**). In the unbound structure, Phe444 side chains in S7b face the center of the ion-conducting pore, erecting a hydrophobic barrier that constricts the pore diameter and prevents ion passage (**Fig. 4C**), similar to the gate observed in ORs^13,14^. When D-fructose binds, these hydrophobic groups swing counter-clockwise away from the pore and are replaced by the polar side chains of Gln443. These rearrangements, accompanied by outward movements of the extracellular ends of the S7b pore helices, result in a widened pore with hydrophilic character (**Fig. 4D**). Gln443 and Phe444 are part of the only signature sequence found in the insect chemoreceptor superfamily (TYhhhhh<u>QF</u>, where h is any hydrophobic amino acid)^23,24^, and mutating either residue to Ala inactivated the receptor (**Fig. 4E,F**). However, mutation of Gln443 to Glu resulted in enhanced activity and increased apparent affinity for D-fructose, suggesting that a more hydrophilic residue at this position favours the activated state of the channel. The hydrophobic to hydrophilic ‘wetting’ transition was also observed in MhOr5^14^ (**Sup. Fig. 7**), suggesting it is likely a common feature of insect chemoreceptor gating.

**Figure 4:**
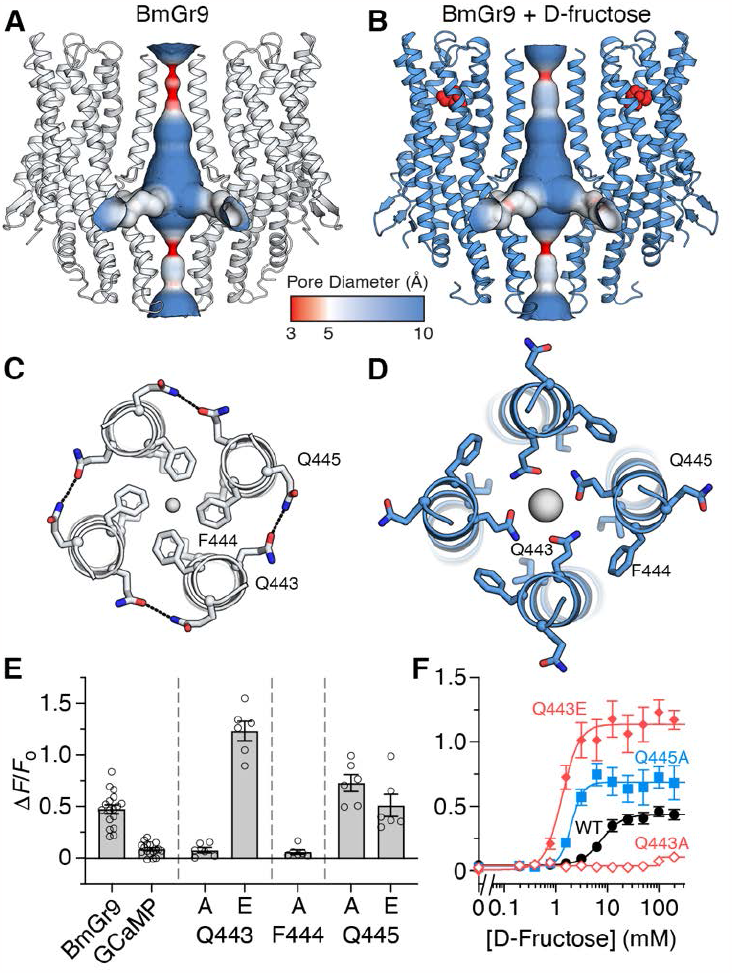
Gating of BmGr9. (**A,B**) The ion permeation pathway of BmGr9, coloured according to pore diameter, in the absence (**A**) and presence (**B**) of D-fructose (red spheres). (**C,D**) Close-up views of the pore helices shown from the top, highlighting key residues in the wetting transition of BmGr9. Hydrogen bonds between Gln443 and Gln445 of adjacent subunits in the closed state are indicated by black dashed lines. Central grey spheres illustrate the narrowest pore diameter, near Phe444 (1.2 Å; **C**) or Gln445 (3.1 Å; **D**). (**E**) Fluorescence changes of BmGr9 and mutants when stimulated with 100 mM D-fructose. (**F**) Dose-response curves of select mutants. Data points are mean points are mean ± s.e.m from *n* = 6–12 experiments.

In the closed state, Gln443 hydrogen bonds with Gln445 from a neighboring subunit (**Fig. 4C**). Eliminating this interaction by mutating Gln445 to Ala created a channel with increased activity and higher apparent affinity compared to the wild-type receptor (**Fig. 4E,F**). Gln445 mutations that retain hydrogen-bonding potential (to Glu or Asn), however, yielded activity similar to wild-type channels (**Sup. Fig. 8**). Thus, without Gln445 to stabilize the position of Gln443 outside of the pore, Gln443 can swing into the pore more easily. This interaction illustrates a possible mechanism for gating cooperativity within the tetramer: ligand binding to one subunit will induce movement of S7b, breaking the Gln443-Gln445 interaction and thus freeing Gln443 of the neighbouring subunit to face the pore. We reassessed the structures of Orco and MhOr5 and identified similar inter-subunit interactions, between Tyr466 and Gln472 in Orco^13^ and Asn469 and Gln467 in MhOr5^14^ (**Sup. Fig. 7**), suggesting S7b-S7b inter-subunit hydrogen-bonded connections are conserved among this chemoreceptor superfamily.

### An aromatic bridge connects pocket and pore

Comparing the bound and unbound structures of BmGr9 reveals a series of helix movements and side chain reorientations that occur upon D-fructose binding. Helices S1, S3, S4, and S6 remain largely fixed, while S2 and S5 move to constrict the pocket around D-fructose (**Fig. 5A**). These conformational changes appear to be driven by specific interactions with D-fructose: Asp99 in S2 moves ∼3 Å to hydrogen bond with the hydroxyl group on C2, and Phe333 in S5 moves ∼2 Å to interact with a second hydrophobic face in D-fructose, formed by hydrogens on C4, C5, and C6. In the absence of a ligand, the vestibule connecting the pocket to the extracellular surface is approximately 8 Å wide, large enough for sugar molecules to access the bottom of the pocket freely (**Fig. 5B**). Upon binding, the tunnel shrinks to 3 Å wide, reducing the pocket volume by about half and tightly enveloping the sugar (**Fig. 5C**).

**Figure 5:**
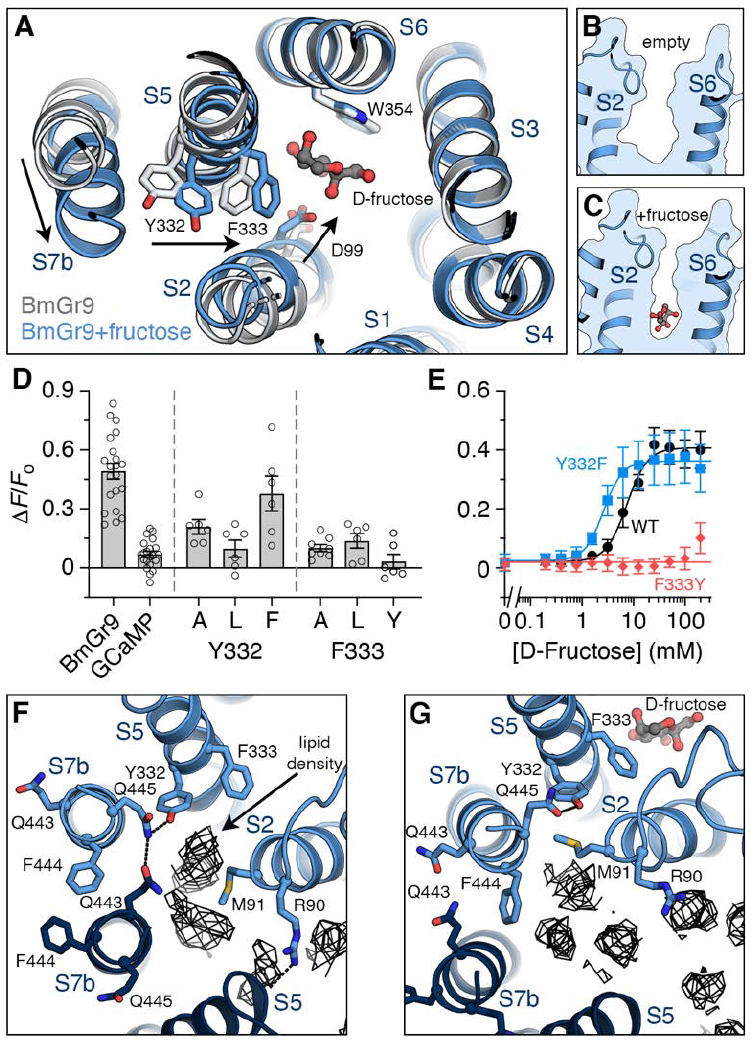
Aromatic bridge connecting D-fructose to channel pore. (**A**) Conformational changes upon binding D-fructose. Arrows indicate movement of key regions in BmGr9 from unbound (grey) to bound (blue) state. (**B,C**) Pocket volume decreases upon binding D-fructose. Outline of the sugar-binding pocket in the absence (**B**) or presence (**C**) of D-fructose. (**D**) Fluorescence changes of BmGr9 and aromatic bridge mutants when stimulated with D-fructose. (**E**) Dose-response of select mutants. (**F,G**) Locations of columnar lipid-like density (wire mesh) reorganize between the unbound (**F**) and bound (**G**) conformations of BmGr9.

Between the pocket and the pore sit Tyr332 and Phe333, which form an ‘aromatic bridge’ on S5 that directly connects D-fructose and residues on the S7b pore helices (**Fig. 5A**). The concerted movement of these residues when D-fructose binds may serve as a switch to open the pore: Phe333 shifts toward D-fructose, pulling the adjacent aromatic residue Tyr332 away from S7b, creating space for the pore helix to move. Tyr332 and Gln445 share a hydrogen bond in both the unbound and bound states, maintaining the direct link between the positions of S5 and S7b. Mutation of Tyr332 to Phe slightly increases the apparent affinity for D-fructose, while to Ala or Leu decreases or eliminates activity, respectively (**Fig. 5D,E**). These varied responses with different amino acids at position 332 suggest that the S5-S7b connection can be fine-tuned to modulate pore opening, independent of specific receptor-ligand interactions within the pocket. By contrast, mutation of Phe333 to Ala, Leu, or Tyr significantly reduced or eliminated activity. This intolerance to modification may reflect the additional need for Phe333 to directly engage the bound sugar.

In the closed state, Tyr332 and Phe333 are exposed to the membrane interior through a gap between the S7b and S2 helices, where lipid-like density is observed in the structure of unbound BmGr9 (**Fig. 5F**). However, when the pore opens, Leu441 in S7b closely associates with Met91 and Val95 in S2, expelling the putative lipid and forming a network of interactions that shield the aromatic bridge from the membrane (**Fig. 5G**). Nearby, Arg90 in S2 reaches across the gap between subunits to interact with C-terminal carbonyls of S5 in the absence of ligand. Movement of S2 towards D-fructose breaks this interaction, increasing the space between subunits and accommodating the shift in lipid positions near the aromatic bridge. This transformation of the membrane-facing surface of BmGr9 between the open and closed states of the receptor raises the possibility that the lipid environment will affect the gating properties of the receptor.

### Mechanism of receptor tuning

Although BmGr9 is only activated by D-fructose, whether other sugars can bind to the receptor is a central matter for determining the origin of its selectivity. To identify other sugars that can potentially bind BmGr9, we computationally docked sweet molecules into the fructose-bound structure of BmGr9. We found that most hexoses similar in size to D-fructose fit well into the binding pocket and make many of the same contacts, yielding similar docking scores (**Sup. Fig. 9**). For example, the lowest energy poses for both D-fructose and L-sorbose are almost identical (**Fig. 6A,B**). Even the change in the C5-hydroxyl position is relatively small, and L-sorbose makes the same number of hydrogen-bonding interactions with the receptor as D-fructose. However, larger sugars, such as the disaccharide sucrose, are too big to fit and their lowest energy poses sit outside the pocket.

**Figure 6:**
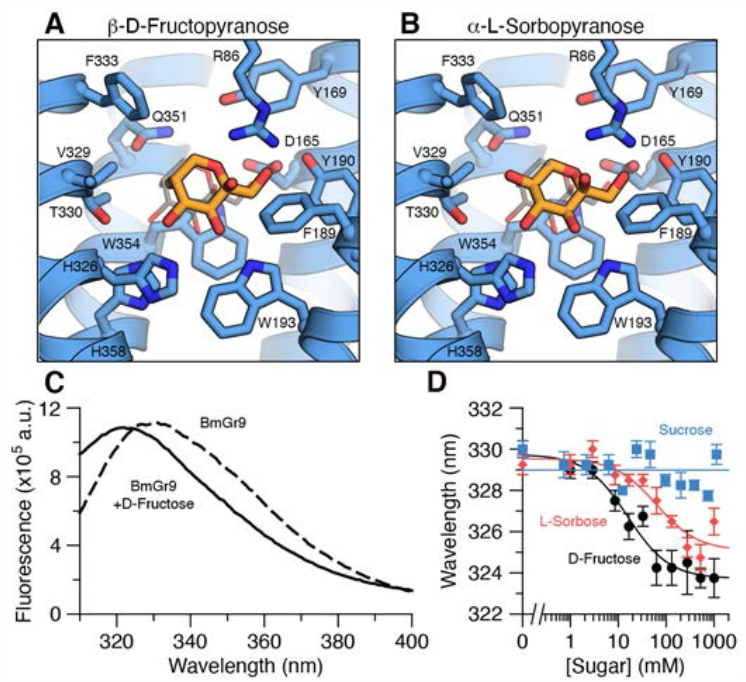
Many hexoses bind BmGr9. (**A,B**) Lowest energy docked poses of β-b-D-fructopyranose (**A**) and α-L-sorbopyranose (**B**) are shown in orange. Thin grey lines indicate the refined position of modelled β-D-fructopyranose. (**C**) Representative tryptophan fluorescence emission spectra of BmGr9 in the absence (dashed line) and presence (solid line) of 100 mM D-fructose. (**D**) Titration of purified BmGr9 with hexoses induces a blue shift in the tryptophan fluorescence emission spectrum while larger sugars do not. The fitted *K*_D_ for D-fructose and L-sorbose are 15.8 (6.4–38.5) mM and 63 (28– 140) mM, respectively (*n* = 4; points are mean ± s.e.m.; fitted 95% confidence intervals are given in parentheses).

To determine experimentally whether sugars with favourable docking scores bind BmGr9, we measured intrinsic tryptophan fluorescence of detergent-solubilized BmGr9. The sugar-binding site in BmGr9 contains two Trps (Trp193 and Trp354), which become less exposed to water once D-fructose is bound. The Trp fluorescence emission spectrum of BmGr9 has a maximum near 330 nm; adding a saturating amount of D-fructose decreases the emission intensity and blue shifts the spectrum to a maximum near 320 nm (**Fig. 6C**), consistent with the Trp residues being buried by D-fructose. The apparent affinity of purified BmGr9 for D-fructose determined using this fluorescence-based assay (*K*_D_ = 16 mM; **Fig. 6D**) is in close agreement with our measurement in cells (8 mM; see **Fig. 1B**) and with previous work on BmGr9^6,16^.

Titration of BmGr9 with several hexoses produces a blue shift in the Trp fluorescence spectrum, similar to that induced by D-fructose (**Fig. 6D**). Consistent with our docking results, L-sorbose binds BmGr9 with an affinity close to that of D-fructose whereas sucrose did not induce a shift in the tryptophan fluorescence spectrum (**Fig. 6D**). As other sugars can bind BmGr9, the precise positioning of amino acids in the pocket is insufficient, by itself, to fully explain the narrow tuning of the receptor.

Since other hexoses are predicted to make many of the same contacts as D-fructose, why do they fail to activate BmGr9? D-Fructose and L-sorbose differ by the orientation of a single hydroxyl group at the C5 position. The structure of BmGr9 bound to D-fructose reveals that the C5-hydroxyl forms a hydrogen bond with Thr330, leaving a hydrophobic patch consisting of aliphatic hydrogens from C4, C5, and C6 to face Phe333 (**Fig. 6A**). In our docked model of L-sorbose, the inverted stereochemistry of C5 places a hydroxyl group in the middle of the hydrophobic patch, likely impeding proper engagement of the aromatic bridge and preventing binding being transduced to pore opening. (**Fig. 6B**).

Our results thus suggest that receptor tuning in BmGr9 arises from two distinct but complementary mechanisms: (1) the stereoelectronic characteristics of the pocket that allow binding, and (2) the engagement of an allosteric process required to activate the channel. Altering contributors to these two elements results in different effects on BmGr9 activity. While mutations within the ligand-binding pocket generally abolished channel activity, many amino acid substitutions along the allosteric pathway were not only tolerated, but created channels that were more active than wild-type BmGr9 (i.e., Y332F, Q443E, and Q445A). Taken together, our data suggest that pocket characteristics have large effects on receptor chemical sensitivity, by modulating which ligands can be recognized, whereas alterations along the allosteric pathway change how binding events are coupled to pore opening, providing a mechanism to tune the selectivity of channels that share similar binding pockets.

## Discussion

Sweet taste receptors serve the essential role to identify necessary nutrients, while also contributing to the pleasurable perception of consuming sweet foods. We have determined the first structures of a eukaryotic sweet taste receptor, offering a unique entry point to investigate the biophysical basis for sweet taste and providing a foundation for understanding how closely related sugars may be discriminated.

BmGr9 is part of an ancient and highly conserved subfamily of nutrient sensors expressed in the brains and mouthparts of insects, which are only activated by D-fructose^6,7^. Sugar specificity in BmGr9 is partly achieved by the arrangement of specific amino acids in the ligand-binding pocket, which creates a set of interactions that precisely match the overall shape and pattern of chemical groups in D-fructose. Residues that line the pocket are highly conserved among fructose-selective GRs, but not in other sugar-sensing GRs, consistent with the chemistry of the ligand binding pocket shaping sugar specificity. Notably, we observed that Trp354 orients the D-fructose molecule within BmGr9 so that its hydroxyl groups can be coordinated by a constellation of polar interactions. Aromatic residues are often present within the predicted binding pockets of GRs and ORs, raising the possibility that they serve a widespread fundamental role in orienting ligands, comparable to other residues that interact with defined chemical groups through ionic or hydrogen bonds^25,26^.

Despite the seemingly specific interactions for D-fructose, our computational docking and experimental binding assays show that several other hexoses can bind BmGr9. Thus, receptor-ligand interactions in the pocket cannot explain the selective activation by only D-fructose if this selectivity is defined by sugar binding alone. We found that D-fructose not only binds BmGr9, but is also unique in its ability to induce a conformational change that is required to open the channel gate, thereby coupling ligand binding to pore opening. The striking specificity of BmGr9 for a single chemical is therefore accomplished by singular complementarity between D-fructose and the aromatic bridge. Other molecules may bind in the pocket, but they are not able to shift the aromatic bridge into an open configuration. The challenge of discriminating between molecules that differ only in the relative positions of a few hydroxyl groups might be too great for pocket structure alone, necessitating an additional layer of chemical selection in BmGr9.

Most sugar-sensing GRs are activated by numerous sweet compounds^10^. BmGr9, therefore, represents an extreme example of specificity within this family. How other GRs achieve broad selectivity is unclear. One possibility is that their binding pockets can interact with many more sugars than BmGr9. This mechanism would resemble that of MhOr5, where the generic binding pocket adapts to accommodate differently shaped odorants^14^. Our work suggests an alternate possibility: that many sugars may bind each GR, but only some can induce the appropriate conformational changes necessary to reach the activation threshold. A likely scenario is that both mechanisms contribute to defining receptor tuning: coarse receptor tuning is derived from the size and chemical characteristics of the pocket, which restricts the set of binding molecules, whereas fine-tuning of receptor activation is achieved through the selective engagement of an allosteric pathway that connects the pocket to the pore. BmGr9 would then occupy one extreme of the tuning spectrum, with broadly tuned chemoreceptors at the other, potentially being activated by almost any molecule able to fit into the ligand-binding pocket.

This two-layer mechanism is likely a general feature of other chemoreceptor families. Inhibition of insect and mammalian ORs by odorants is prevalent^27,28^, indicating that many molecules bind but do not activate these receptors. This inhibition provides an important feature of the combinatorial coding of odour mixtures, suggesting that the coupling between ligand-binding and receptor activation may be a point of evolutionary selection that can tune the activity of receptors with similar binding pockets. Continued investigation into chemoreception by diverse receptors will help us explore the relationship between amino acid sequence, specificity, and receptor tuning— ultimately revealing how families of receptors decipher the chemical world.

## Data availability

The final cryo-EM maps have been deposited in the Electron Microscopy Data Bank under accession numbers EMD-42629 (fructose-bound) and EMD-42628 (unbound). The final models have been deposited in the Protein Data Bank under accession numbers 8UVU (fructose-bound) and 8UVT (unbound). Source activity and binding data are provided with this paper. For all other data requests, please contact J.A.B.

## Acknowledgments

We thank J. Carlson, M. Lemmon, B. Noro, V. Ruta, and members of the Butterwick lab for comments on this manuscript and advice throughout the course of this project. We thank K. Yedlin for early experiments with BmGr9 and greatly appreciate the support from M. Llaguno and S. Wu for cryo-EM grid screening and data collection, B. Evans for assistance with cryo-EM data analysis software, and L. Abriola for activity measurements. J.V.G was supported by the Brazilian Federal Agency for Support and Evaluation of Graduate Education (CAPES). This work was funded by generous grants from the Bill and Melinda Gates Foundation and the Whitehall Foundation (to J.A.B.).

## Author contributions

J.V.G. purified BmGr9, collected and analyzed cryo-EM data, and performed GCaMP activity measurements. S.S-B. performed docking computations. J.V.G, C.C.C., and M.S. measured tryptophan fluorescence. M.C. performed the initial screening of sugar-sensing GRs and preliminary expression tests. J.A.B. supervised all aspects of the research. J.V.G and J.A.B. wrote the manuscript with input from all authors.

## Materials & Methods

### Expression and purification of BmGr9

A synthetic construct consisting of residues Pro2– Ser449 (the native C-terminus) of *B. mori* Gr9 (Genbank Accession EU769120.1) was cloned into a pEG BacMam vector (Addgene plasmid #160451; from E. Gouaux)^29^ following an N-terminal Strep-tag II^30^, superfolder GFP^31^ and an HRV 3C protease site. Baculovirus containing the BmGr9 coding sequence was created in Sf9 cells (ATCC CRL-1711). HEK293S GnTI^−^ cells (ATCC CRL-3022) were grown in suspension at 37 °C in Freestyle 293 medium (Gibco) supplemented with 2% (v/v) fetal bovine serum (FBS; Gibco) and 1% (v/v) GlutaMAX (Gibco) with 8% (v/v) carbon dioxide until they reached a density of ∼3×10^6^ cells/ml and then transduced with baculovirus at a multiplicity of infection of ∼1. After 12 h, 10 mM sodium butyrate (Sigma-Aldrich) was added to the medium, and the temperature was reduced to 30 °C. The cells were harvested ∼48 h later by centrifugation and washed once in phosphate-buffered saline (pH 7.5; Gibco). Cell pellets were frozen in liquid nitrogen and stored at – 80 °C until needed.

Cell pellets were thawed on ice and resuspended in 20 mL of lysis buffer per gram of cells. Lysis buffer was composed of 50 mM HEPES/NaOH (pH 7.5), 375 mM NaCl, ∼10 µg/mL DNase I, 1 µg/mL leupeptin, 1 µg/mL aprotinin, 1 µg/mL pepstatin A, 1 mM phenylmethylsulfonyl fluoride (all from Sigma-Aldrich). BmGr9 was extracted by adding 1% (w/v) n-dodecyl-ß-D-maltoside (DDM; Anatrace) with 0.2% (w/v) cholesterol hemi-succinate (CHS; Sigma-Aldrich) for 2 h at 4°C. The mixture was clarified by centrifugation at 80,000*g* and the supernatant was added to 0.4 mL StrepTactin Sepharose resin (Cytiva) per gram of cells and rotated at 4 °C for 1 h. The resin was collected, washed with 10 column volumes (cv.) of 20 mM HEPES/NaOH (pH 7.5), 150 mM NaCl (HBS) with 0.02% (w/v) DDM, 0.004% (w/v) CHS, and then with 10 cv. of HBS with 0.05% (w/v) digitonin (Sigma-Aldrich). BmGr9 was eluted by adding 2.5 mM desthiobiotin (Sigma-Aldrich) to the digitonin buffer.

The StrepII–GFP tag was cleaved by HRV 3C protease (Novagen) added at 10 U/mg of BmGr9 overnight at 4 °C. BmGr9 concentration was calculated from its absorbance at 280 nm assuming an extinction coefficient (ε_280_) of 45.8 mM^−1^ cm^−1^ (calculated by ProtParam^32^). BmGr9 was then concentrated in a centrifugal tube (Amicon Ultra-4; 100 kDa cutoff) and injected onto a Superose 6 Increase column (Cytiva) previously equilibrated with HBS with 0.05% (w/v) digitonin. For the fructose-bound sample, 0.5 M D-fructose (Sigma-Aldrich) was included in the buffer.

### Cryo-EM sample preparation and data collection

Peak fractions containing purified BmGr9 were concentrated to 4.7 mg/mL (unbound sample) or 2.9 mg/mL (fructose-bound sample). Cryo-EM grids were frozen using a Vitrobot Mark IV (FEI) as follows: 3 µL of the concentrated sample was applied to a glow-discharged Quantifoil R1.2/1.3 holey carbon 400 mesh gold grid, blotted for 2.5–4 s in >90% humidity at room temperature, and plunge frozen in liquid ethane cooled by liquid nitrogen. Grids were screened for ice thickeness and particle distribution using a Glacios (200 kV; Thermo Scientific) in the Yale School of Medicine Center for Cellular and Molecular Imaging.

Cryo-EM data were recorded on a Titan Krios (300 kV; FEI) in the Yale West Campus Cryo-EM Core, equipped with a Gatan K3 Summit camera and imaging filter. SerialEM^33^ was used for automated data collection. Movies were collected at a nominal magnification of 81,000× in super-resolution mode resulting in a calibrated pixel size of 0.534 Å/pixel, with a defocus range of approximately –1.0 to –3.0 µm. Fifty frames were recorded over 10 s of exposure at a dose rate of 1.67 electrons per Å^2^ per frame. Movie frames were aligned and binned over 2×2 pixels using MotionCor2^34^ and the contrast transfer function (CTF) parameters for each motion-corrected image were estimated using CTFFIND4.1^35^.

An initial set of ∼2000 BmGr9 particles were manually picked and submitted to 2D class average to create references for auto-picking. For the unbound BmGr9 sample, 1,179,559 particles from 5770 micrographs were extracted, into 384×384-pixel boxes, binned over 3×3 pixels, and subjected to further 2D classification using RELION-3.1^36^. After removing junk particles, the remaining 446,673 particles were re-extracted without binning and used to build an initial 3D model in RELION-3.1. The model was further refined with C4 symmetry imposed, ultimately reaching 3.8 Å resolution without masking. The particles were then imported into CryoSPARC (Structura Biotechnology)^37^ and underwent a new round of 2D classification. After subsequent rounds of non-uniform refinement (with C4 symmetry), local CTF refinement, and further 2D classification, we obtained a map with a nominal resolution of 2.9 Å, estimated using the Fourier shell correlation (FSC) = 0.143 cutoff^38^.

A similar approach was used for the fructose-bound BmGr9. 1,760,600 particles from 7035 micrographs were auto-picked in RELION-3.1. After initial 2D classification, 1,157,505 particles were extracted and moved to CryoSPARC for further processing. We used 307,715 particles to build the final map with a 3.0 Å resolution. The density images in **Fig. 1C** and **Sup. Fig. 3D,H** were created using UCSF ChimeraX^39^.

### Cryo-EM data analysis and model building

Both maps were of sufficient quality for de novo atomic model building. A poly-alanine model for BmGr9 was built in Coot^40^, and subsequent amino acid assignments were made based on side-chain densities. The models were refined using real-space refinement implemented in PHENIX41 for five macro-cycles with four-fold non-crystallographic symmetry and secondary structure restraints applied. The lowest scoring docking poses (see below) were used as the starting positions to refine the structures of D-fructose conformers. Ligand restraints were obtained using electronic Ligand Builder and optimization Workbench (eLBOW)42 implemented in PHENIX.

There was no apparent density for the intracellular loop between S4-S5, hence both final models lack this region. Model statistics were obtained using MolProbity43. Models were validated by randomly displacing the atoms in the original model by 0.5 Å, and refining the resulting model against half maps and full map. Images of the model in Figs. 1-6 and Sup. Fig. 7 were created using PyMOL44 and in Sup. Figs. 4,5,9 using UCSF ChimeraX. Fischer and Haworth projections of sugars were made using ChemDraw 22.2 (PerkinElmer).

### Structural analysis

Residues at subunit interfaces or the binding pocket were identified using PyMOL as any residue within 5 Å of a neighboring subunit or the ligand, respectively. Hydrogen bonds shown in Fig. 2 were identified using PyMOL (polar contacts).

The pore diameters along the central axis and lateral conduits were calculated using the program HOLE45 with 0.5 Å step sizes (Fig. 4A-D). Two calculations were performed: one along the central four-fold axis (central pore) and another between subunits near the cytosolic membrane interface (lateral conduits). The pores overlapped in the central vestibule.

### Computational Docking

Molecular docking was performed with the unbound and fructose-bound models and a ligand set of various sweet compounds using the molecular docking software AutoDock Vina19,20 with the Vinardo46 scoring function. As some carbohydrates assume different conformations in solution, all anomeric forms were considered. Cubical grids of different sizes and locations in and around the protein were generated using the grid feature of AutoDock Tools to determine the best docking space within the protein. In the end. a 4.500 A3 cubical grid was centered in the observed binding pocket with x- and y-axis lengths of 15 Å and a z-axis length of 20 Å. The structures were prepared in AutoDock Tools by adding any missing atoms and charges on residues assuming a pH of 7.4. Compound structures (SDF files) were downloaded from PubChem or ZINC databases and prepared for docking by conversion into PDBQT files using OpenBabel47.

### Cell-based GCaMP calcium flux assay

The cell-based GCaMP assay was based on a previously described method used to study insect ORs^13^. BmGr9 variants were cloned into a modified pME18s vector with no fluorescent tag. Each transfection condition contained 0.5 µg of a plasmid encoding GCaMP6s (Addgene plasmid #40753; from D. Kim & GENIE Project) and 1.5 µg of the plasmid encoding the BmGr9 variant, in 250 µl OptiMEM (Gibco). These were mixed with a solution containing 7 µl Lipofectamine 3000 (Invitrogen) in 250 µl OptiMEM, followed by a 15-min incubation at room temperature.

HEK293 cells were maintained in high-glucose DMEM (Gibco), supplemented with 10% (v/v) FBS and 1% (v/v) GlutaMAX, and 5% (v/v) carbon dioxide at 37 °C. Cells were detached using trypsin and resuspended at a concentration of 10^6^ cells/ml. Cells were mixed with the transfection mixture and added to a 384-well plate with a clear bottom (Grenier). Four to six hours later, a 16-port vacuum manifold on low vacuum was used to remove the transfection medium, which was replaced by fresh FluoroBrite DMEM (Gibco) supplemented with 10% (v/v) FBS and 1% (v/v) GlutaMAX. Twenty-four hours later, this medium was replaced with 50 µl reading buffer (20 mM HEPES/NaOH (pH 7.4), 0.1× Hank’s Balanced Salt Solution (Gibco), 3 mM Na_2_CO_3_, and 5 mM CaCl_2_) in each well.

The fluorescence emission at 515–575 nm, with excitation at 475–495 nm, was continuously read by a FLIPR Tetra System (Molecular Devices) at the Yale Center for Molecular Discovery. The exposure time was set to 0.5 s, excitation intensity at 100%, and camera gain adjusted according to the baseline signal for each plate. After 30 s of baseline recording, 25 µl of tastant solution was added to the cells and read for 8 min. All solutions were pre-warmed to 37 °C. All sweet compound titrations were made using eleven ligand concentrations for each transfection condition in sequential dilutions of 2, alongside controls wells of reading buffer only. Ligands were dissolved in the reading buffer at 600 mM, then diluted with reading buffer to the highest final-well concentration of 200 mM. Due to their lower solubility, the sweeteners aspartame and saccharin had the highest final-well concentration of 10 mM.

Each plate contained negative control wells transfected with GCaMP6s alone and exposed to tastants. Additionally, each plate included BmGr9 with its cognate ligand D-fructose as a positive control to monitor plate-to-plate variation in transfection efficiency and cell count. Each ligand concentration was applied to two technical replicates, averaged, and considered a single biological replicate. The baseline fluorescence (Fo) was calculated as the average fluorescence of the 30 s before any addition to the plate. The maximum signal was reached 50-70 s after tastant addition, and the average fluorescence signal in that period (F) was used for further calculations. F-Fo/Fo for each concentration was calculated to account for well-to-well variability, In the sugar panel in Fig. 1A. percent activity was calculated as the difference between fluorescence change evoked by adding 100 niM of sweet compound (or 10 mM of aspartame or saccharin) between cells transfected with BmGr9 plus GCaMP and control cells transfected with only GCaMP, divided by the difference observed when treated with D-fructose (multiplied by 100 to yield a percentage). For all experiments, GraphPad Prism 10 was used to fit four-parameter Hill equation to dose-response data, from which £Cso and Hill coefficients were extracted.

#### Tryptophan fluorescence measurements

BmGr9 was expressed and purified as described above. Sugar solutions were made in the same buffer used for the Superose 6 column. 100 µL of BmGr9 (at 1 µM) was added to a quartz cuvette, and the tryptophan fluorescence was measured by a PTI fluorometer using the Felix software (version 1.42b. PTI). The excitation wavelength was set to 295 mn with a bandwidth of 20 inn. Fluorescence emission was measured at 300 to 400 mn. with a 1 nm step size. After adding each ligand, solutions were mixed gently, followed by fluorescence measurements after 1 min. All curves were corrected for dilution. The same ligand concentrations were added to 25 µM n-acetyl-L-tryptophanamide (NATA) in 0.05% digitonin to account for potential compound inner filter effects. The observed fluorescence quenching in the NATA experiments was subtracted from BmGr9 measurements for every ligand. All data were processed using Excel (Microsoft) and the maximum emission wavelengths analyzed using GraphPad Prism 10.

### Sequence alignments

All sequence alignments were visualized and plotted using JalView48. All protein sequences were obtained in the UniProtKB or GenBank databases.

### AlphaFold tetramer & structural comparison

We used ColabFold49 to predict the structure of BmGi9 homotetramer. Four identical primary sequences of BmGr9 were added to the AlphaFold prediction tool in Chimera X and submitted with default parameters to Google Colab servers. Images showing model comparisons were generated using PyMOL and UCSF ChimeraX.

## Supplementary Figures & Tables

**Supplementary Figure 1:**
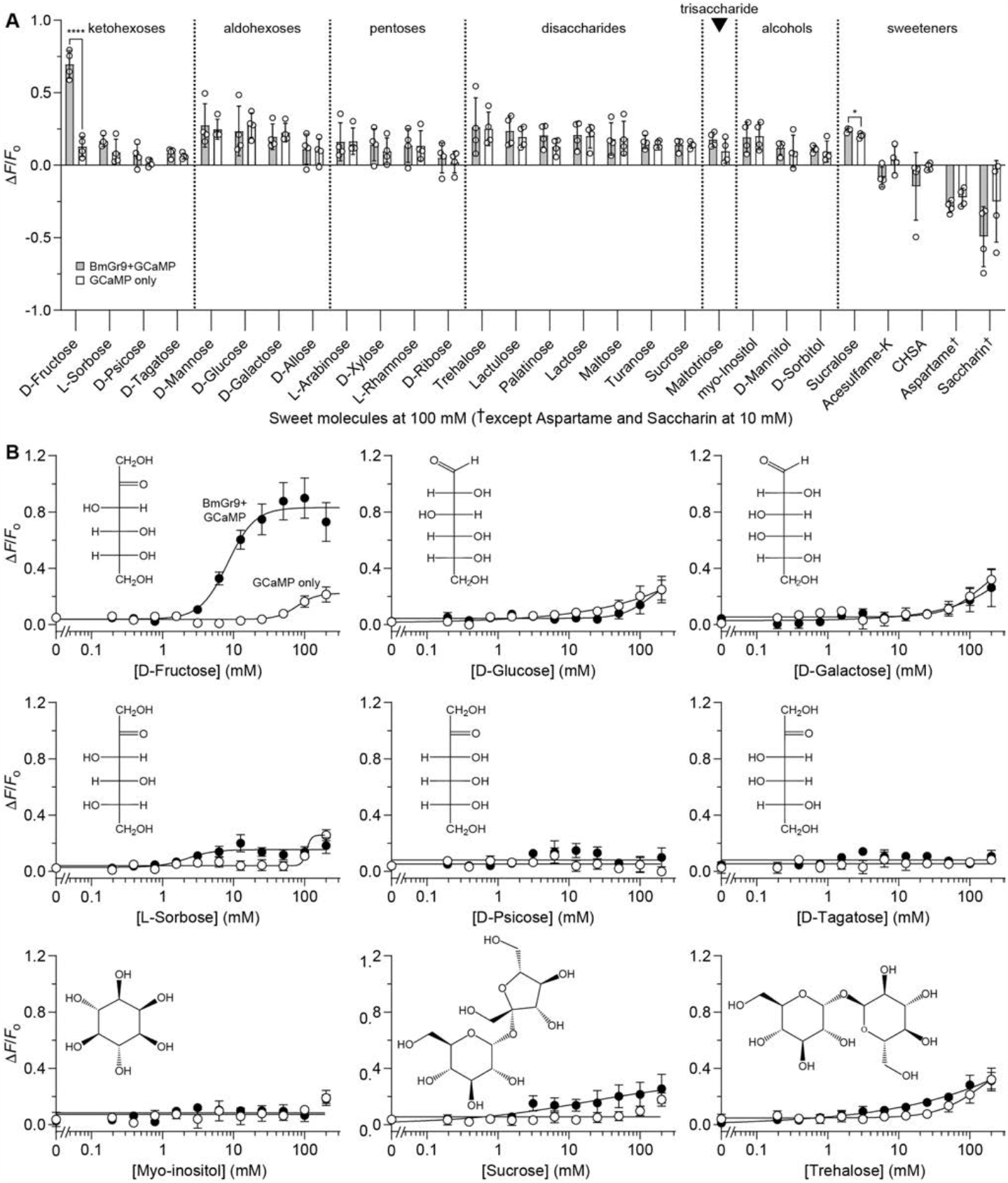
BmGr9 is narrowly tuned to D-fructose. (**A**) Change in fluorescence upon the addition of sweet compounds (at 100 mM, except for aspartame and saccharin, which were at 10 mM) in HEK293 cells transfected with BmGr9 plus GCaMP (grey bars) or with GCaMP only (white bars). Bars are mean ± s.e.m with individual replicates shown as open circles (*n* = 4). Only D-fructose (*P* < 0.0001) and sucralose (*P* < 0.05) additions yielded significantly different activity with BmGr9. (**B**) Dose-response of fluorescence changes of HEK293 cells transfected with BmGr9 and GCaMP (closed circles) or GCaMP only (open circles) when titrated with select sugars from (**A**) (*n* = 4–8, points are mean ± s.e.m.). Insets show Fischer projections of hexoses and perspective projections of myo-inositol, sucrose, and trehalose.

**Supplementary Figure 2:**
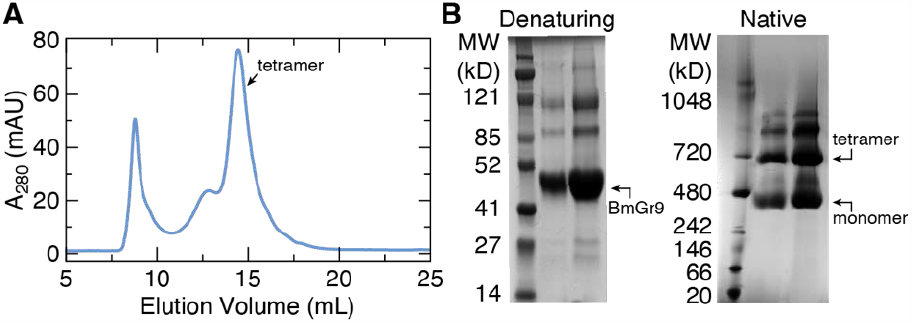
Purification of BmGr9. (**A**) Superose 6 elution profile of purified BmGr9. The majority of the protein elutes as a tetramer. (**B**) Coomassie staining of denaturing and native gels confirm BmGr9 is a homotetramer with an effective molecular weight of approximately 700 kDa (including detergent micelle), similar to Orco^13^. Molecular weight markers are labeled for each gel.

**Supplementary Figure 3:**
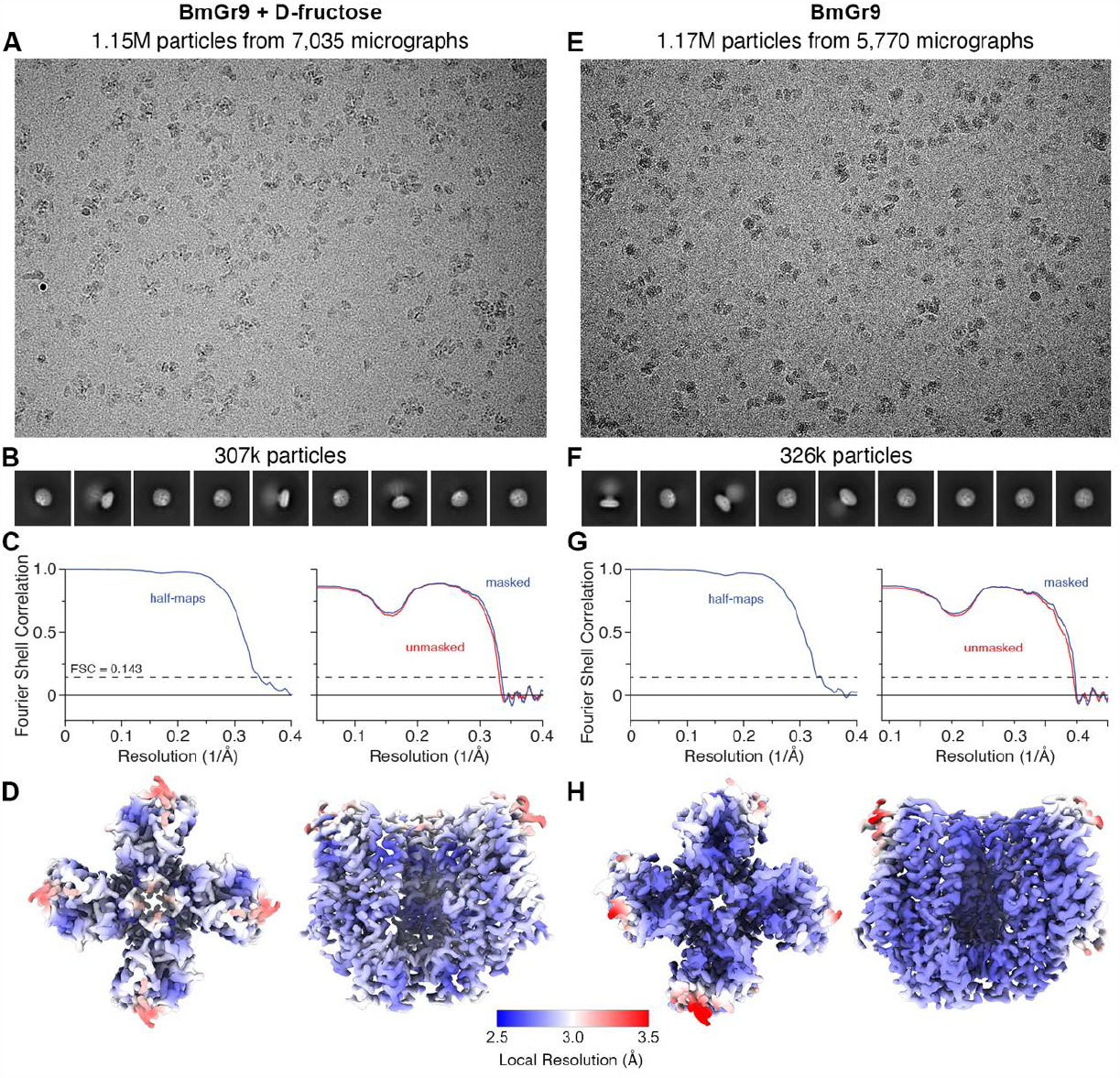
Cryo-EM workflow. (**A**) A representative motion-corrected micrograph showing the distribution of fructose-bound BmGr9 single particles. The numbers of micrographs and auto-picked particles are shown. (**B**) Example two-dimensional class averages of particles selected for further processing. (**C**) Fourier shell correlation (FSC) curves for the final cryo-EM density maps. Half-map FSC (with tight mask) (left), model-map FSC curves (right). The horizontal dashed line represents the FSC = 0.143 cutoff value. (**D**) Local resolution of fructose-bound BmGr9 density map. (**E-H**) Equivalent data for unbound BmGr9.

**Supplementary Figure 4:**
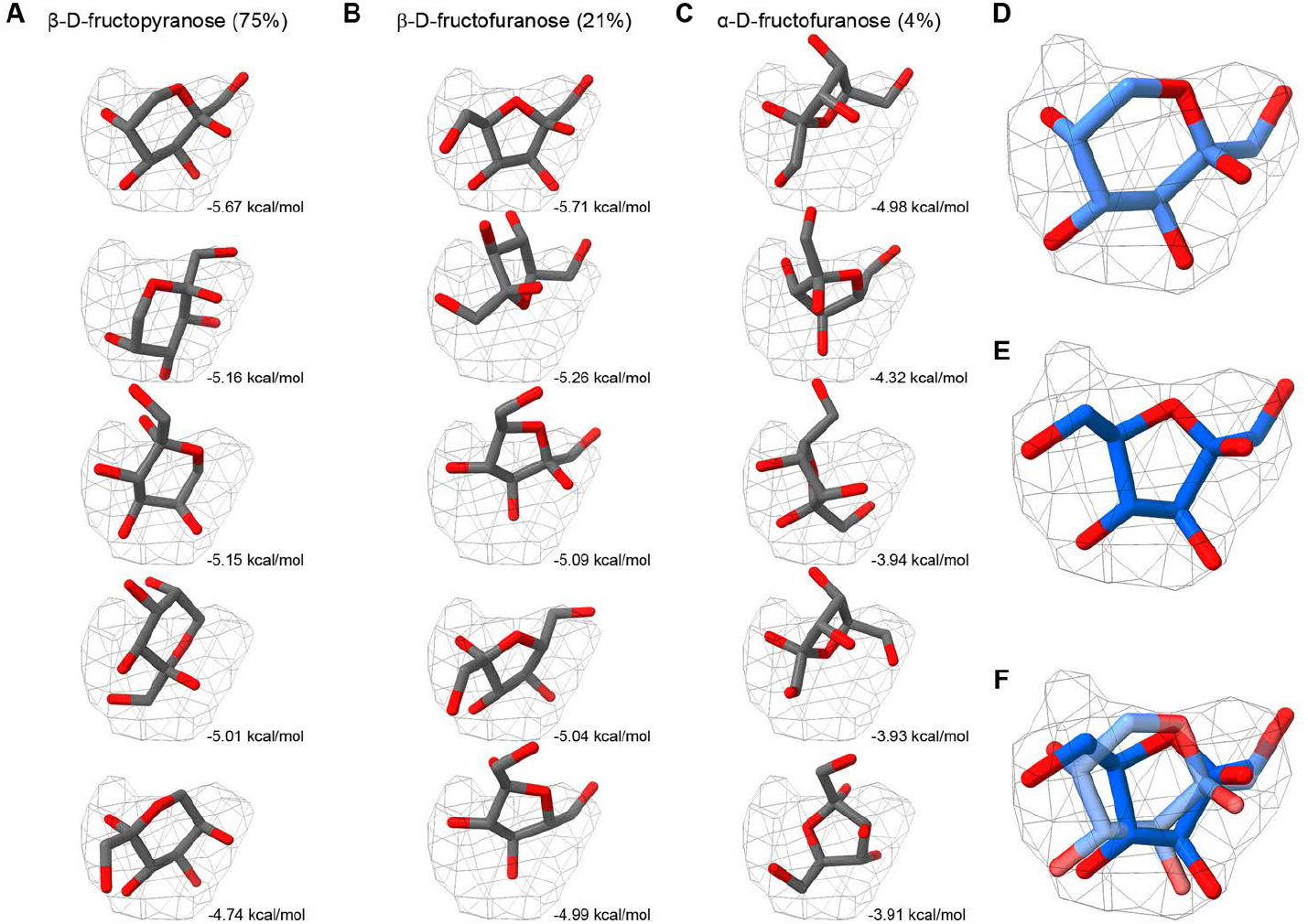
Computational docking of D-fructose anomers. (**A-C**) Five lowest energy poses for β-D-fructopyranose (**A**), β-D-fructofuranose (**B**), and α-D-fructofuranose (**C**), and their respective energy scores. Experimental ligand density is shown as a grey mesh. The equilibrium composition in solution at room temperature is shown in parentheses^17,18^. (**D,E**) The final positions of of β-D-fructopyranose (**D**, light blue) and β-D-fructofuranose (**E**, dark blue) after real-space refinement. (**F**) Superimposition of refined β-D-fructopyranose and β-D-fructofuranose positions.

**Supplementary Figure 5:**
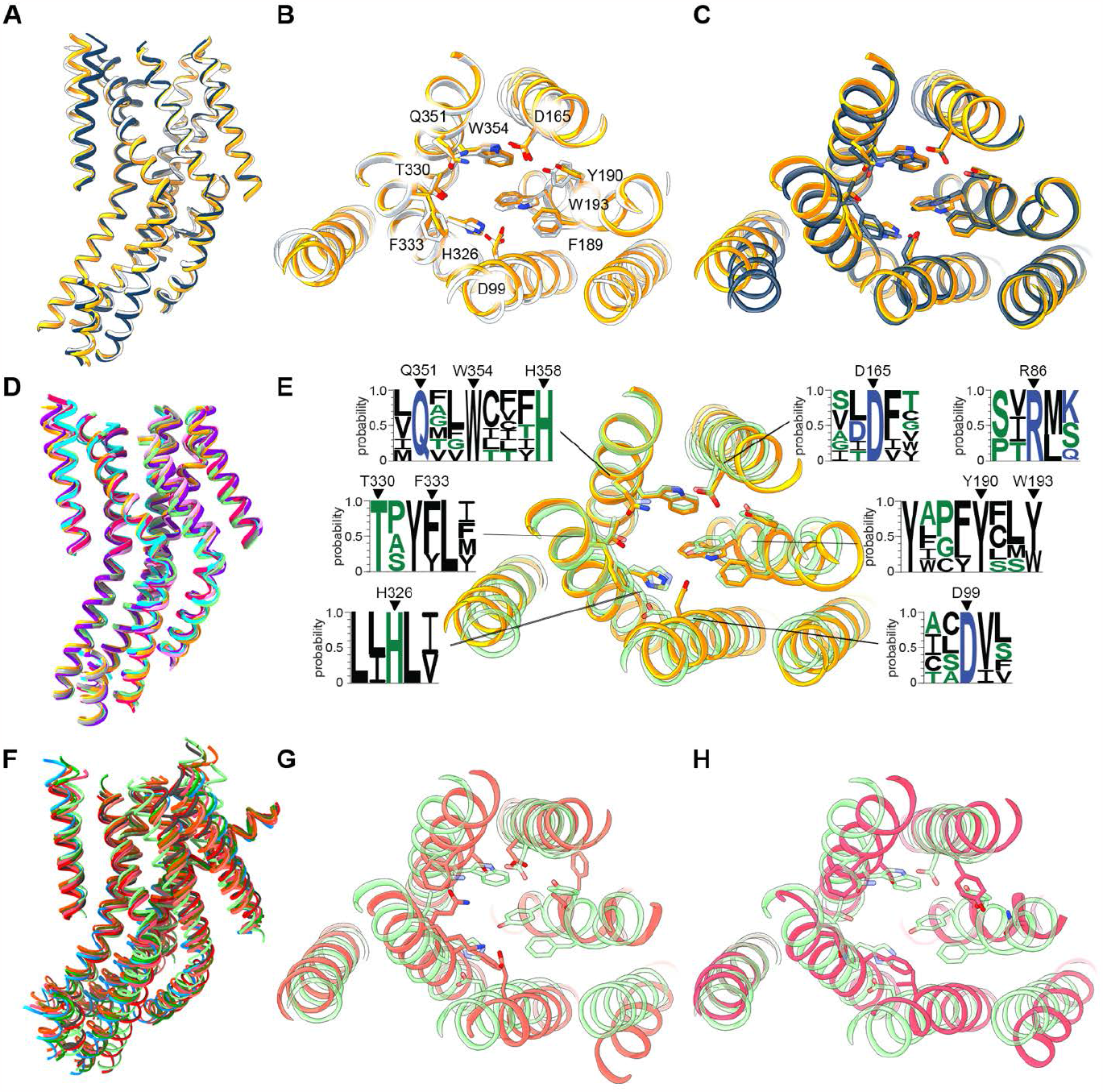
Structure conservation among sugar-sensing GRs. (**A-C**) Superposition of fructose-bound BmGr9 (blue), unbound BmGr9 (white), and BmGr9 predicted using AlphaFold2 (yellow). (**B,C**) Top views of the ligand-binding pocket with residues shown (and labelled in (**B**)). (**D**) Aligned AlphaFold2 models of known Gr43a-like receptors: BmGr9 (yellow); BmGr10 (purple); DmGr43a (light green); *Anopheles gambiae* Gr25 (cyan); *Helicoverpa armigera* Gr9 (grey); *Apis mellifera* Gr3 (light pink); and *Trichogramma chillonis* Gr43a (dark pink). (**E**) Comparison of AlphaFold2 models of BmGr9 and DmGr43a, with pocket residues shown. Logo representation of amino acid conservation among the Gr43a-like receptors in (**D**). Residues that interact with D-fructose are highlighted. (**F**) AlphaFold2 models of *D. melanogaster* sugar GRs: DmGr5a (light red); DmGr43a (light green); DmGr61a (light blue); DmGr64a (magenta); DmGr64b (dark green); DmGr64c (red); DmGr64d (dark grey); DmGr64e (orange); and DmGr64f (gold). (**G,H**) Comparison of AlphaFold2 models of DmGr43a (light green) and (**G**) DmGr5a (light red) or (**H**) DmGr64a (magenta) with pocket residues shown. No residues in the pocket are conserved between these receptors. In all images, loops are hidden for clarity.

**Supplementary Figure 6:**
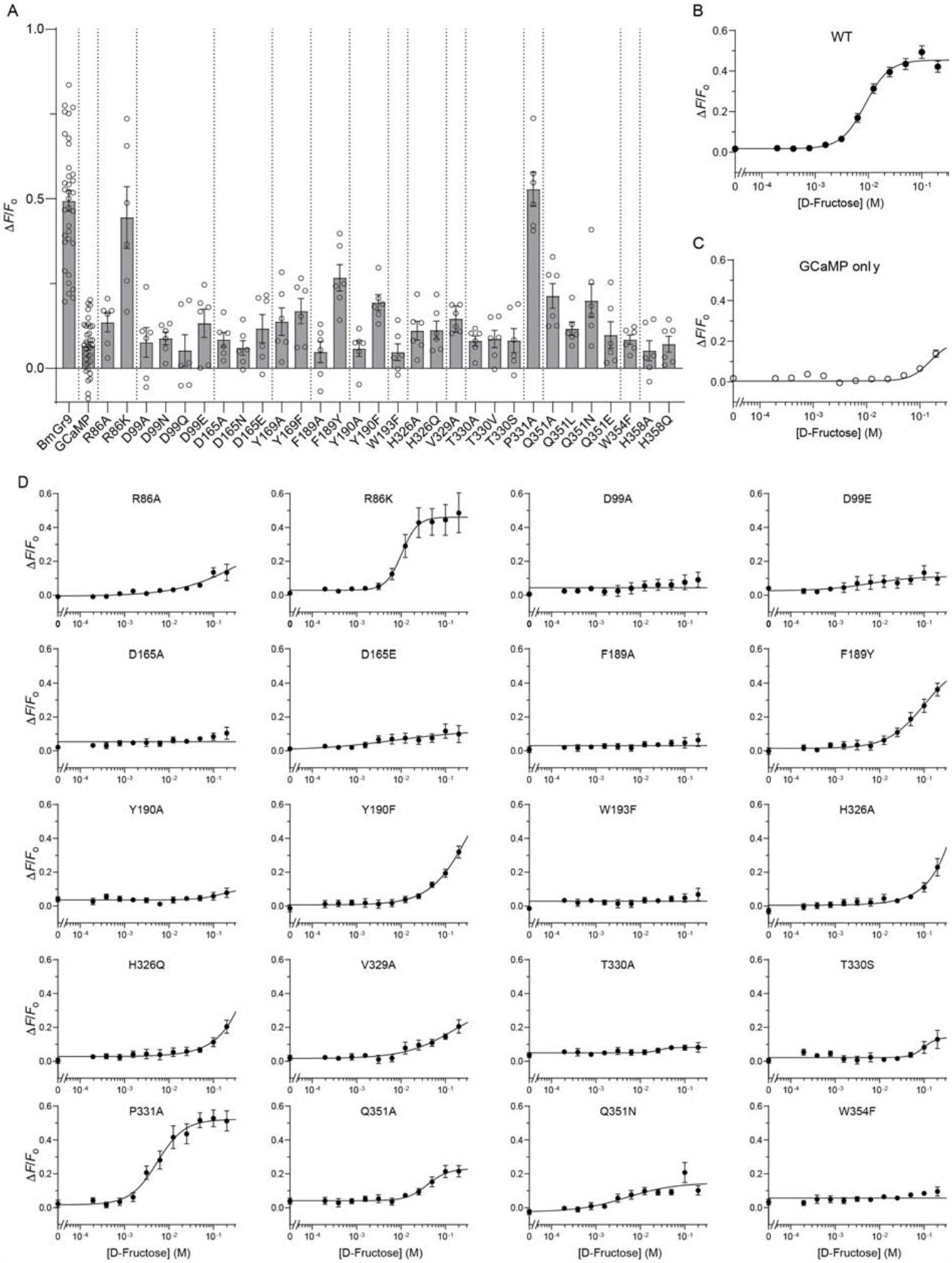
Mutational analysis of the sugar-binding pocket in BmGr9. (**A**) Fluorescence changes of BmGr9 and mutants when stimulated with 100 mM D-fructose. Bars are mean ± s.e.m with individual replicates shown as open circles. Wild-type BmGr9 and GCaMP-only controls are indicated. (**B-D**) D-Fructose dose-response curves of HEK293 cells transfected with wild-type BmGr9 plus GCaMP (**B**), GCaMP only (**C**), and select mutants (**D**). Data points are mean ± s.e.m from *n* = 6–12 experiments.

**Supplementary Figure 7:**
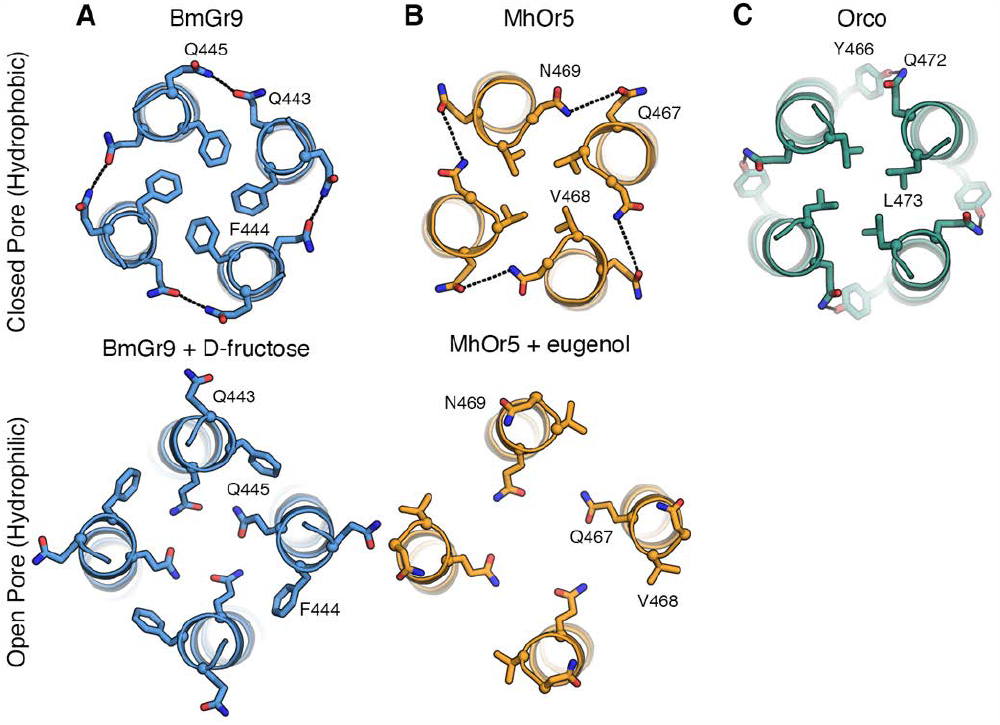
Conserved gating in insect chemoreceptors. Comparison of pore helices in BmGr9 (**A**), MhOr5 (**B**), and Orco (**C**) in the close (top) and open configurations (bottom). In the closed states, hydrophobic resisdues line the channel gates, which are replaced by hydrophilic residues in the open states. Intersubunit interactions are shown as dashed lines. PDB codes are: 7LIC (MhOr5), 7LID (MhOr5 bound to eugenol), and 6C70 (Orco).

**Supplementary Figure 8:**
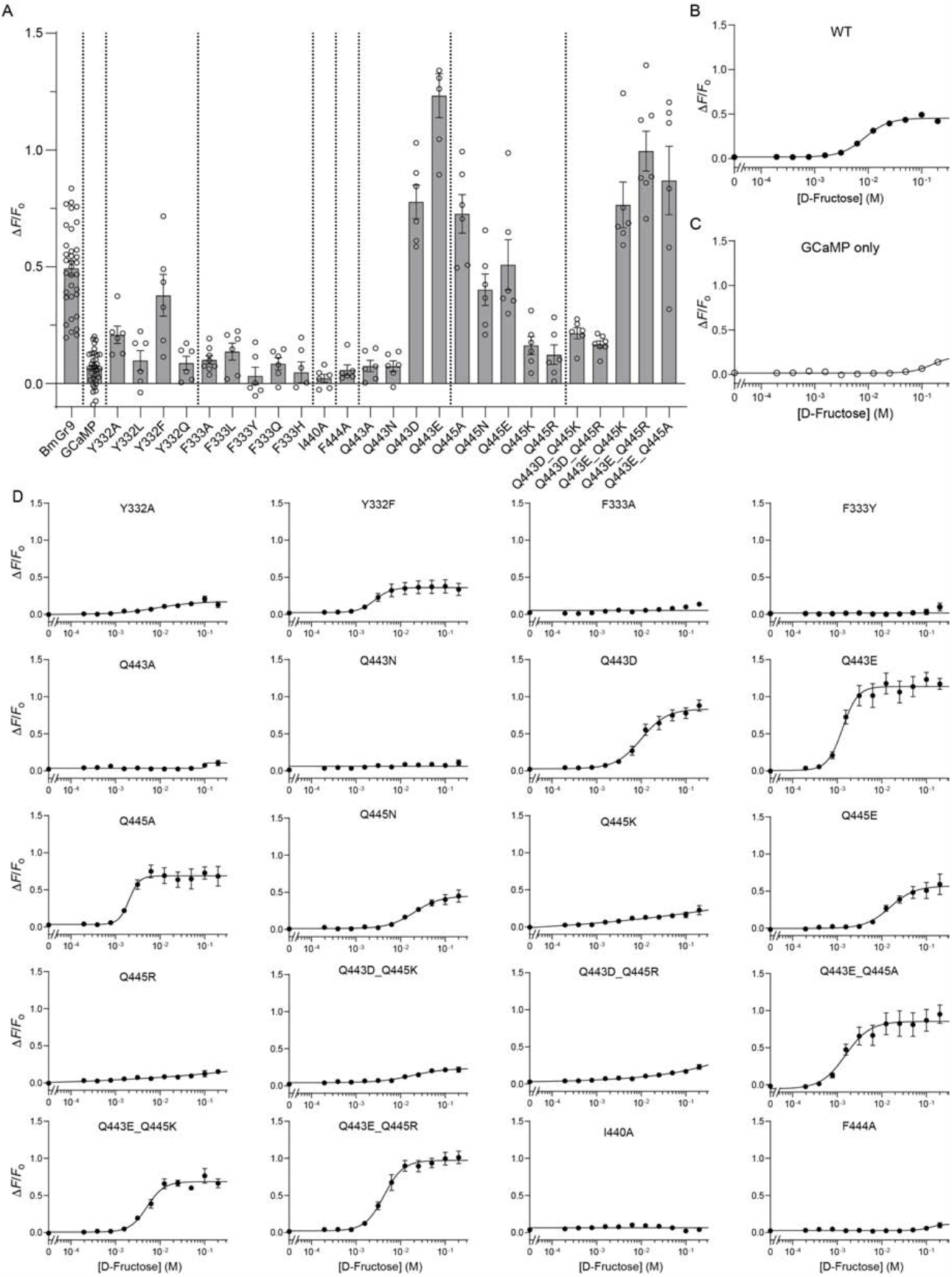
Mutational analysis of BmGr9 gating. (**A**) Fluorescence changes of BmGr9 and mutants when stimulated with 100 mM D-fructose. (**B-D**) D-Fructose dose-response curves of HEK293 cells transfected with (**B**) wild-type BmGr9 plus GCaMP, (**C**) GCaMP only, and (**D**) select mutants. Data in (**B,C**) are the same as in Sup. Fig. 6, but with y-axis adjusted for scale. Data points are mean points are mean ± s.e.m from *n* = 6–12 experiments.

**Supplementary Figure 9:**
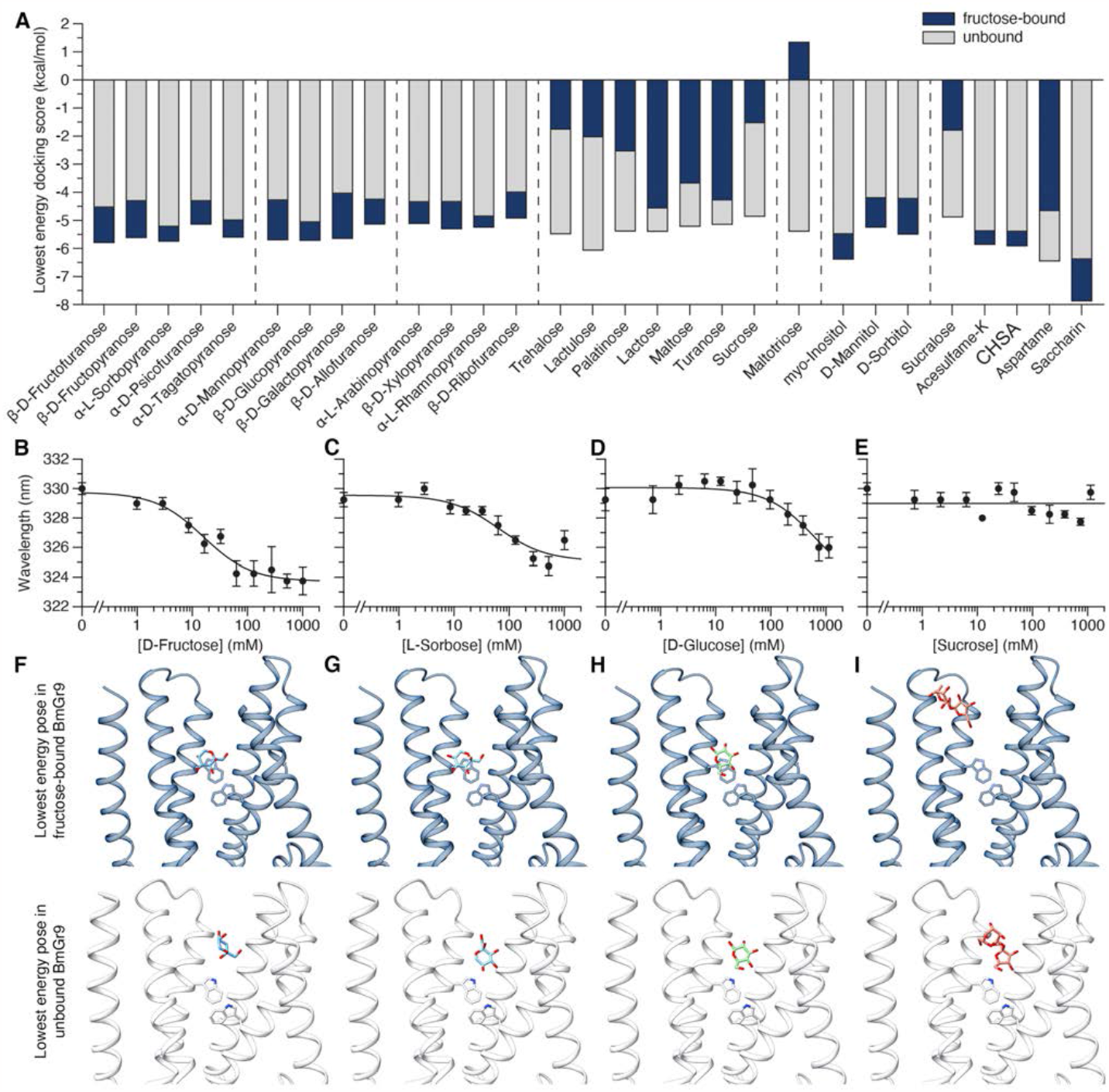
Many sweet molecules bind to BmGr9. (**A**) Superimposed bars of the docking scores (kcal/mol) of sweet molecules in fructose-bound BmGr9 (blue) and apo-BmGr9 (grey). Only predominant anomers in solution were selected, except for D-fructose. (**B-E**) Wavelength of the maximum tryptophan fluorescence emission spectrum of BmGr9 when titrated with D-fructose (**B**), L-sorbose (**C**), D-glucose (**D**), and sucrose (**E**). (**F-I**) Lowest energy poses for docked β-D-fructopyranose (**F**), α-L-sorbopyranose (**G**), β-D-glucopyranose (**H**), and sucrose (**I**) into the fructose-bound structure of BmGr9 (blue, top) or unbound structure (white, bottom).

**Supplementary Table 1:** Cryo-EM data collection, refinement and model statistics.

|  | BmGr9<br>EMDB-42628<br>PDB 8UVT | BmGr9 + Fructose<br>EMDB-42629<br>PDB 8UVU |
| --- | --- | --- |
| **Data collection and processing** |  |  |
| Magnification | 81,000× | 81,000× |
| Voltage (kV) | 300 | 300 |
| Electron exposure (e-/Å<sup>2</sup>) | 1.67 | 1.67 |
| Defocus range (μm) | −0.8 to −2.5 | −0.8 to −2.5 |
| Pixel size (Å) | 1.068 | 1.068 |
| Symmetry imposed | C4 | C4 |
| Initial particle images (no.) | 1,179,559 | 1,157,505 |
| Final particle images (no.) | 326,748 | 307,715 |
| Map resolution (Å) | 2.92 | 3.05 |
| FSC threshold | 0.143 | 0.143 |
| Map resolution range (Å) | 2.3–8.0 | 2.5–9.0 |
| **Refinement** |  |  |
| Initial model used (PDB code) | n/a | n/a |
| Model resolution (Å) | 4.0 | 4.0 |
| FSC threshold | 0.143 | 0.143 |
| Map sharpening *B* factor (Å<sup>2</sup>) | −147.5 | −151.5 |
| Model composition |  |  |
| Non-hydrogen atoms | 12148 | 11928 |
| Protein residues | 1548 | 1516 |
| Ligands | 0 | 4 |
| *B* factors (Å<sup>2</sup>) |  |  |
| Protein | 22.04 | 28.14 |
| Ligand | n/a | 17.69 |
| R.m.s. deviations |  |  |
| Bond lengths (Å) | 0.002 | 0.002 |
| Bond angles (°) | 0.494 | 0.389 |
| Validation |  |  |
| MolProbity score | 1.82 | 1.60 |
| Clashscore | 9.28 | 5.28 |
| Poor rotamers (%) | 0 | 0 |
| Ramachandran plot |  |  |
| Favoured (%) | 95.25 | 95.47 |
| Allowed (%) | 4.75 | 4.53 |
| Disallowed (%) | 0 | 0 |

n/a, not applicable

**Supplementary Table 2:** BmGr9 and mutant activation by D-fructose.

| | *N* | $F_{\max}$ | Hill Slope | $EC_{50}$ (mM) |
| --- | --- | --- | --- | --- |
| WT | 34 | 0.45 (0.43 to 0.48) | 1.94 (1.48 to 2.58) | 8.71 (7.40 to 10.29) |
| GCaMP | 34 | ~0.20 | ~2.37 | ~154 |
| R86A | 6 |  | no activity |  |
| R86K | 6 | 0.46 (0.39 to 0.56) | ~2.45 | 10.4 (6.7 to 17.2) |
| D99A | 6 |  | no activity |  |
| D99N | 6 |  | no activity |  |
| D99E | 6 |  | no activity |  |
| D99Q | 6 |  | no activity |  |
| D165A | 6 |  | no activity |  |
| D165N | 6 |  | no activity |  |
| D165E | 6 |  | no activity |  |
| F189A | 6 |  | no activity |  |
| F189Y | 6 | ~0.53 | 1.05 (0.44 to 2.15) | ~0.10 |
| Y190A | 6 |  | no activity |  |
| Y190F | 6 | ~0.88 | 0.98 (0.56 to 2.07) | ~363 |
| W193F | 6 |  | no activity |  |
| H326A | 6 | ~70 | 0.95 (0.51 to 3.76) | ~87190 |
| H326Q | 6 | ~41 | ~0.92 | ~73630 |
| V329A | 6 | ~0.35 | 0.79 (0.28 to 2.08) | ~153 |
| V329T | 6 | ~2.82 | ~0.64 | ~19810 |
| T330A | 6 |  | no activity |  |
| T330V | 6 | ~0.13 | ~6.22 | ~98 |
| T330S | 6 | ~0.14 | ~3.03 | ~100 |
| P331A | 6 | 0.52 (0.47 to 0.60) | 1.43 (0.87 to 2.39) | 5.5 (3.7 to 8.5) |
| Q351A | 6 | 0.23 (0.18 to 0.65) | 1.89 (0.60 to 5.26) | 3.9 (2.3 to 593) |
| Q351N | 6 | ~0.17 | ~0.71 | ~8.0 |
| Q351L | 6 | ~0.10 | ~1.71 | ~25 |
| Q351E | 6 |  | no activity |  |
| W354F | 6 |  | no activity |  |
| H358A | 6 | ~0.15 | ~1.06 | ~365 |
| H358Q | 6 | ~0.14 | ~1.43 | ~113 |
| F333A | 6 |  | no activity |  |
| F333Y | 6 |  | no activity |  |
| F333L | 6 |  | no activity |  |
| F333Q | 6 |  | no activity |  |
| F333H | 6 |  | no activity |  |
| Y332A | 6 | 0.17 (0.14 to 0.23) | 0.99 (0.37 to 2.17) | 8.3 (3.5 to 42) |
| Y332F | 6 | 0.36 (0.31 to 0.43) | ~2.36 | 2.6 (1.3 to 5.0) |
| Y332L | 6 |  | no activity |  |
| Y332Q | 6 |  | no activity |  |
| Q443A | 6 |  | no activity |  |
| Q443E | 6 | 1.14 (1.05 to 1.23) | ~2.57 | 1.32 (0.99 to 1.75) |
| Q443N | 6 |  | no activity |  |
| Q443D | 6 | 0.83 (0.75 to 0.96) | 1.52 (0.94 to 2.57) | 10.0 (7.4 to 14.8) |
| Q445A | 6 | 0.69 (0.62 to 0.75) | ~3.78 | 2.01 (1.15 to 2.71) |
| Q445E | 6 | 0.57 (0.47 to 0.85) | 1.56 (0.68 to 3.75) | 15.3 (9.8 to 42.2) |
| Q445N | 6 | 0.45 (0.38 to 0.62) | 1.49 (0.79 to 2.76) | 20.3 (13.3 to 44.4) |
| Q445K | 6 |  | no activity |  |
| Q445R | 6 |  | no activity |  |
| Q443D_Q445K | 6 | 0.24 (0.20 to 2.72) | 1.18 (0.24 to 2.87) | ~18 |
| Q443D_Q445R | 6 | ~3.13 | 0.31 (0.13 to 1.13) | ~1181000 |
| Q443E_Q445K | 6 | 0.69 (0.64 to 0.74) | 2.28 (1.51 to 3.62) | 5.10 (4.07 to 6.33) |
| Q443E_Q445R | 6 | 0.95 (0.88 to 1.03) | 2.02 (1.27 to 3.33) | 4.07 (3.14 to 5.31) |
| Q443E_Q445A | 6 | 0.85 (0.75 to 1.01) | 1.58 (0.69 to 6.42) | 1.53 (0.88 to 2.82) |
| I440A | 6 |  | no activity |  |
| F444A | 6 |  | no activity |  |
| I335A | 6 |  | no activity |  |

The number of biological replicates (*N*), and fitted values for the maximum fluorescence ( $F_{\max}$ ), Hill slope, and concentration of half-maximal activation ( $EC_{50}$ ) are listed; 95% confidence intervals are given in parentheses. Parameters for which 95% confidence intervals could not be accurately determined are indicated by a tilde (~)

